# Tracking the Metabolic Fate of Exogenous Arachidonic Acid in Ferroptosis Using Dual-Isotope Labeling Lipidomics

**DOI:** 10.1101/2023.05.28.542640

**Authors:** Noelle Reimers, Quynh Do, Rutan Zhang, Angela Guo, Ryan Ostrander, Alyson Shoji, Chau Vuong, Libin Xu

## Abstract

Lipid metabolism is implicated in a variety of diseases, including cancer, cell death, and inflammation, but lipidomics has proven to be challenging due to the vast structural diversity over a narrow range of mass and polarity of lipids. Isotope labeling is often used in metabolomics studies to follow the metabolism of exogenously added labeled compounds because they can be differentiated from endogenous compounds by the mass shift associated with the label. The application of isotope labeling to lipidomics has also been explored as a method to track the metabolism of lipids in various disease states. However, it can be difficult to differentiate a single isotopically labeled lipid from the rest of the lipidome due to the variety of endogenous lipids present over the same mass range. Here we report the development of a dual-isotope deuterium labeling method to track the metabolic fate of exogenous polyunsaturated fatty acids, e.g, arachidonic acid (AA), in the context of ferroptosis using hydrophilic interaction-ion mobility-mass spectrometry (HILIC-IM-MS). Ferroptosis is a type of cell death that is dependent on lipid peroxidation. The use of two isotope labels rather than one enables the identification of labeled species by a signature doublet peak in the resulting mass spectra. A Python-based software, D-Tracer, was developed to efficiently extract metabolites with dual-isotope labels. The labeled species were then identified with *Lipydomics* based on their retention times, collision cross section, and *m/z* values. Changes in exogenous AA incorporation in the absence and presence of a ferroptosis inducer were elucidated.

**Table of Contents:** 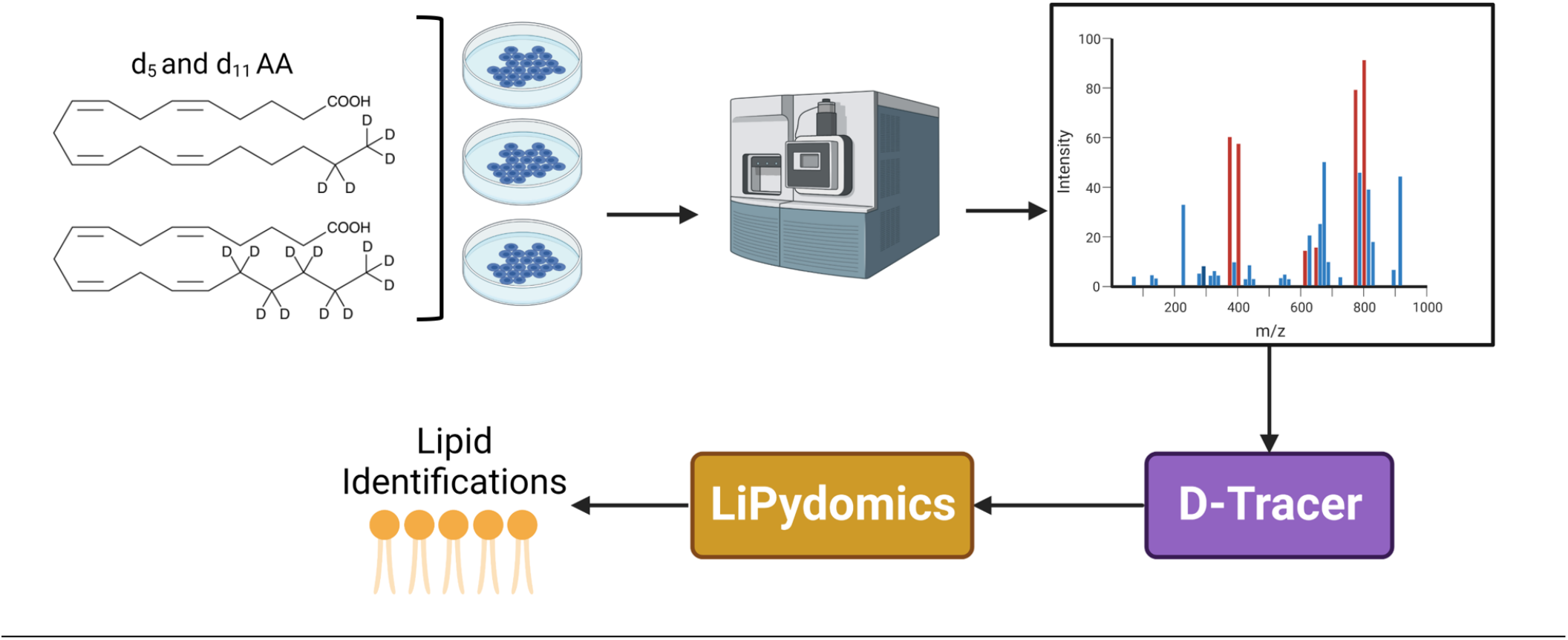

## INTRODUCTION

The rise of genomics, proteomics, and metabolomics over the last 20 years has brought a wealth of information to the field of biochemistry and greatly advanced our understanding of human health and diseases. Lipidomics is a branch of metabolomics that studies lipids and their metabolites within a biological system.^1^ Lipidomics has revealed the many roles of lipids beyond simple energy storage molecules, including as signaling molecules in inflammation and the immune response, metabolic regulation, and cell death.^2^ Altered lipid metabolism is a hallmark of many types of cancer,^3^ as cancer cells often change their lipid metabolism to keep up with their need for energy, alter signaling pathways, and evade cell death.^4,5^ Altered lipid metabolism has also been implicated in cardiac dysfunction,^6^ liver disease,^7^ and neurodegeneration.^8^ Therefore, the advancement of the lipidomics field could yield important insights into these diseases and others.

In recent years, altered lipid metabolism and increased lipid peroxidation were found to play a critical role in the iron-dependent cell death form, ferroptosis.^9–12^ Polyunsaturated fatty acids (PUFAs) are crucial to the process of lipid peroxidation because the hydrogen atoms at the bis-allylic position between two double bonds are especially labile and subject to hydrogen atom abstraction by a radical (Figure 1).^13^ Arachidonic acid (AA) is a biologically relevant omega-6 PUFA with 20 carbon atoms and 4 double bonds at the 5,8,11, and 14 positions. It is found at a concentration of approximately 400 µM in human plasma.^14^ Previous work has shown that AA treatment can sensitize cells to ferroptotic death, enhancing the lethality of canonical ferroptosis inducer RSL3 in cancer cells.^15^ RSL3 inhibits the lipid peroxide-reducing enzyme glutathione peroxidase 4.^10,12^ However, the exact species and biochemical events that lead to peroxide buildup and eventual cell death are unclear.

**Figure 1.**
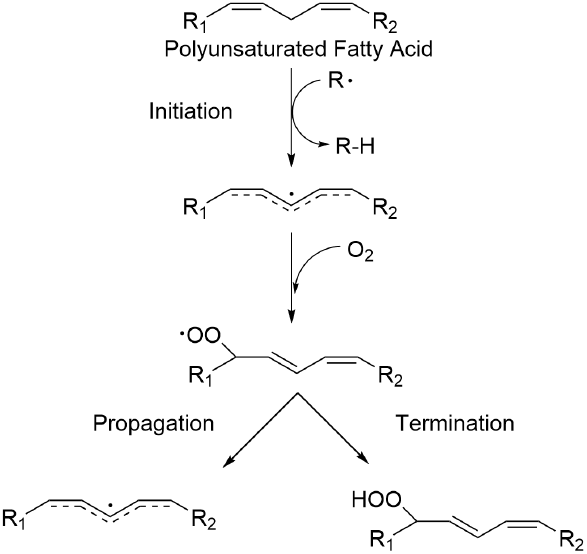
Scheme of free radical peroxidation of PUFAs via the hydrogen atom transfer (HAT) mechanism.

Despite its biological importance, lipidomics has several inherent difficulties that hinder its advancement. There are upwards of 47,000 observed and computational lipids in the Lipid MAPs database, which continues to expand.^16^ Most of these cellular lipids occur in a relatively narrow mass range, from 500 -1200 Daltons. Within this range, there are many isomeric and isobaric lipids that are impossible to identify by mass alone. These traits make the separation, resolution, and identification of lipids quite challenging. Mass spectrometry is the primary tool used in lipidomics due to its resolving power, throughput, and customizability.^1^ The use of chromatography and ion mobility, along with soft ionization techniques such as electrospray ionization (ESI), has greatly improved our ability to resolve many of these compounds.^17^

Most lipids within a cell share the same structural pattern of two or three fatty acid tails attached to a polar or semi-polar headgroup. The headgroup structures have enough variation in their polarity that they can be separated using hydrophilic interaction liquid chromatography (HILIC).^18^ In addition, lipid ions can adopt different conformations while in the gas phase. Depending on the headgroup and degree and location of double bonds within the fatty acid tails, a lipid may be more or less compact in space.^19–21^ Ion mobility (IM) spectrometry can measure the apparent surface area, *i*.*e*., collision cross section (CCS), of an ion as it traverses an inert background gas.^22^ CCS values are shown to be highly reproducible across different IM platforms ^23–25^ and have been used to enhance the confidence of lipid identification.^26,38,40^ While these advances have significantly improved the utility of lipidomics, some challenges remain.

One of these challenges is the ability to trace lipid metabolites, as is commonly done in other areas of metabolomics.^27 13^C and deuterium labeling can be used to trace the metabolism of many compounds, including lipids,^28–30^ amino acids,^31^ and xenobiotics.^32,33^ When a compound is labeled with ^13^C or deuterium, it results in an increase in mass that differentiates it from the endogenous, unlabeled, analog. This mass shift can be detected with analytical methods including mass spectrometry and nuclear magnetic resonance and can be used to identify the compounds into which the molecule is metabolized.^27^ Applying this tool to lipidomics has been challenging because the mass shift associated with the isotopic label is often not enough to differentiate it from other lipids in the lipidome since most lipids occur in a narrow mass range.^34^ It is likely that labeled peaks could be missed or incorrectly identified because they may be indistinguishable from other lipids in the same sample.

To overcome this challenge, we developed a dual-isotope labeling technique that uses an equimolar mix of two different deuterated fatty acids (referred to as dAA). These fatty acids are incorporated into the lipidome equally by the cells, resulting in mass spectra with a signature doublet or triplet peak. These signature spectra, along with the same retention time and CCS values, can be used to recognize the features associated with the metabolites of the labeled fatty acid with confidence. We then developed a Python software, D-Tracer, to automatically pick out labeled metabolite pairs and correct the isotope shift to generate the *m/z* values of the corresponding non-labeled lipids. We integrated this program with our previously developed Python package, *Lipydomics*, to identify the mass-corrected lipids using three-dimensional data that includes *m/z*, chromatographic retention time (RT), and CCS values.^35^ Subsequently, we applied this method to characterize the metabolic fate of labeled AA in HT-1080 endothelial cancer cells in the context of ferroptosis.

## EXPERIMENTAL

### Materials

Mass spectrometry-related materials were previously described. ^26,36^ Briefly, HPLC grade solvents (water, acetonitrile, methylene chloride, chloroform, and methanol) were purchased from Thermo Fisher Scientific. Optima LC/MS grade ammonium acetate and sodium chloride were also obtained from Thermo Fisher. d_5_- and d_11_-AA were purchased from Cayman Chemical (9000477 and 10006758, respectively)

### Lipid internal standards

PE (15:0/15:0), Cer (d18:1/17:0), PG (15:0/15:0), PC (15:0/15:0), PA (12:0/12:0), PS (12:0/12:0), SM (d18:1/17:0), LysoPC(15:0/0:0), LysoPE ((13:0/0:0), GluCer(d18:1/17:0) and d_7_-PI(15:0/18:1) were purchased from Avanti Polar Lipids. DG (13:0/13:0) and TG (15:0/15:0/15:0) were purchased from NuChek. Table S1 lists the catalog numbers of all internal standards used. All standards were dissolved in LC-MS grade methanol at a concentration of 10 mg/mL. To make the internal standard mix, the dissolved internal standards were diluted in DMSO such that each standard was at a final concentration of 240 µM.

### Cell Culture

HT-1080 cells (ATCC) were cultured in DMEM high glucose media containing 10% fetal bovine serum and 1% penicillin-streptomycin (Gibco). 5.0 × 10^5^ cells were plated in 10 cm dishes and incubated at 37 C. 24 hours later, the media was exchanged to media containing 80 µM of each deuterated arachidonic acid, 0.03 µM RSL3, or both. The cells were harvested after another 24 hours of incubation. To harvest the cells, cell culture dishes were first placed on ice. To retain any dead cell debris, the media was collected into labeled glass centrifuge tubes and centrifuged at 1000 RPM for 5 minutes. The media was then aspirated, leaving behind a pellet of cell debris. The remaining cells were then rinsed with PBS, scraped off the plates, and added to the corresponding centrifuge tube. Another round of centrifugation afforded the final cell pellets.

### Lipid Extraction

Lipid extraction from treated cells took place as previously described.^36^ Briefly, cells were collected as described above. The cell pellet was resuspended in 300 µL of ice-cold PBS and sonicated for 30 minutes to lyse. Protein levels were quantified using the Bio-Rad BSA Protein Quantification assay and a BioTek Synergy Microplate Reader. 1 mL of ice-cold 0.9% (w/v) NaCl aqueous solution, 4 mL of ice-cold Folch (2:1 v/v chloroform/methanol), and 12 µL of the 240 µM internal standard mix were added to each tube. The tubes were vortexed and centrifuged at 1000 RPM for 5 minutes. Using a glass Pasteur pipet with a mechanical pipet pump, the resulting lower organic layer was recovered from each tube and placed into a fresh tube. The solution was dried using a speed vacuum concentrator (Thermo Fisher Savant SpeedVac). The dried extract was reconstituted in 300 µL of methylene chloride and transferred to HPLC screw-top vials. Immediately prior to MS analysis, 25 µL of each sample was transferred to an HPLC vial with an insert and dried under nitrogen, then reconstituted in 100 µL of acetonitrile:methanol (2:1) for injection.

### HILIC-IM-MS Method

The samples were run on a Waters Synapt XS ion mobility-QTOF mass spectrometer with ESI as previously described.^36^ The instrument was calibrated for CCS using PC and PE standards before use.^37^ Sodium formate was used for mass calibration in both positive and negative modes. HILIC A mobile phase is 95% acetonitrile, 5% water, and 5 mM ammonium acetate while HILIC B mobile phase is 50% acetonitrile, 50% water, and 5 mM ammonium acetate. The stationary phase used was a HILIC Phenomenex Kinetex 2.1 x 100 mm, 1.7 µm, column. Leucine enkephalin was used as a lock mass and CCS calibrant in both modes. The injection volume was 5 µL and solvent flow rate was 0.5 mL/min. The TriWave settings were as described in Table S2.^36^

The samples were injected in order with a blank and pooled sample between each treatment group. Each run lasted 12 minutes starting with 100% HILIC A and dropping to 70% HILIC A by 8 minutes.

### Data Analysis

Data analysis was carried out using Waters Progenesis QI, EZ Info, and two in-house Python programs. First, Progenesis is used to align spectra, perform lock-mass calibration, and pick peaks. Treatment groups are compared to each other using ANOVA. Compounds that are significantly (corrected *P*-value < 0.05) different between dAA-treated and untreated groups are exported to EZinfo for OPLS-DA analysis. Significantly increased features in the treatment group relative to controls are selected from the resulting S-plot and exported as a .csv file.

To find labeled lipid species, we developed a Python software, D-Tracer, with a web interface. The software searches for species that have an RT within 0.01 minutes, a CCS within 3%, and an *m/z* separated by (D2 – D1) x the mass of deuterium. D1 and D2 are user inputs that can be customized. In our case, D2 is 11, and D1 is 5. The mass of the deuterium is then subtracted from each identified species, and the output .csv contains the original *m/z*, adjusted *m/z*, RT, CCS, and normalized intensities for each lipid. The number of samples is also a user input that can be customized. The software has been tested on peaklists containing up to 6,000 compounds and can complete analysis of these large datasets in less than 5 minutes. The source code is available on GitHub under the name D-Tracer (https://github.com/nreimers99/d-tracer).

To identify lipid species, we used LiPydomics, which is a Python software developed by our group.^35^ Identifications are further confirmed using the LipidMaps database. To quantify each lipid species, the intensities of individual lipid species were compared with an internal standard from the same class (Table S1).

## RESULTS AND DISCUSSION

### Development of D-Tracer

There are existing programs commonly used to analyze lipidomics data and identify lipids based on their *m/z*, RT, and CCS values, including LipidIMMS,^38^ MS-Dial 4,^39,40^ and *Lipydomics*.^35^ However, these programs could not be used to identify isotope-labeled lipids due to the isotopic mass shift from deuterium incorporation. To identify the lipids using established databases, the mass of deuterium-containing lipids must first be adjusted to the mass of the corresponding non-labeled lipid. Some programs can analyze single-isotope labeling experiments,^41–43^ but, to our knowledge, no program is currently available for dual-isotope labeling lipidomics. To address this, we developed a web-based Python program called D-Tracer to find and adjust the mass of deuterated lipids. The program was designed to be intuitive and accessible to those without prior coding experience. The simple web version can be accessed by a url (https://nreimers99-d-tracer-original-d-tracerapplication-xji04u.streamlit.app/; see Supplementary Information for detailed instructions), whereas the full version can be used in the terminal. The program can analyze dual isotope deuterium labeling experiments with any degree of deuteration from 0 to 80 and any .csv peaklist if it contains RT, CCS, and *m/z* columns. A schematic of D-Tracer is shown in Figure 2A.

**Figure 2.**
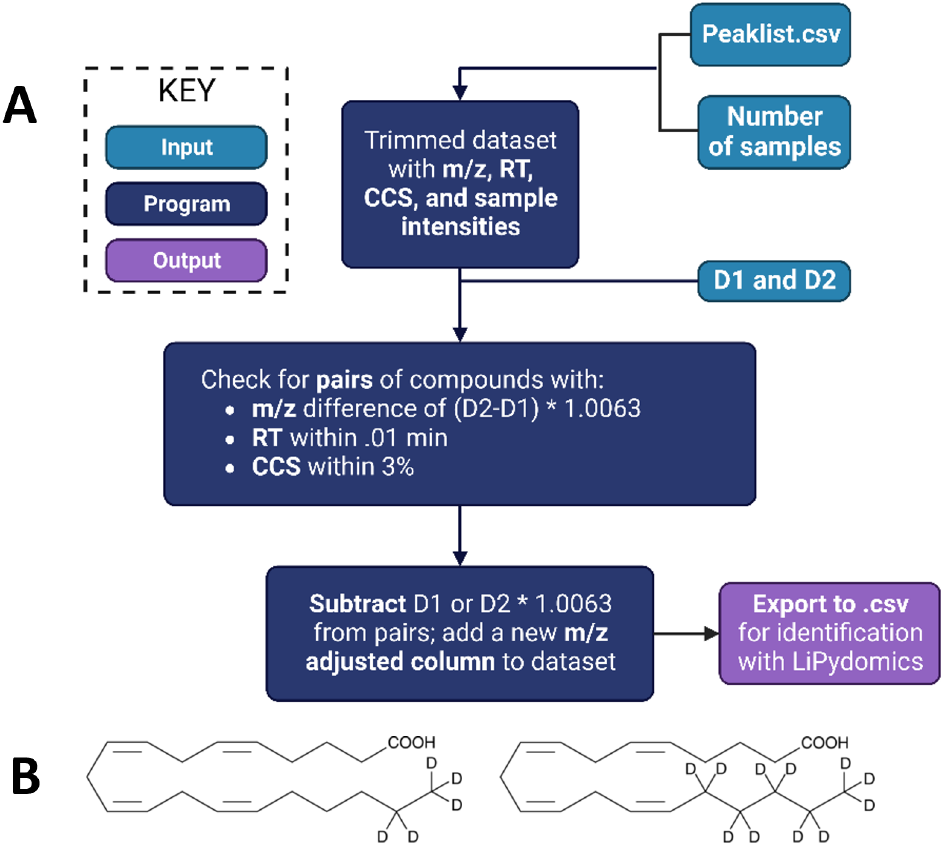
Schematic of the Python-based software D-Tracer. User inputs into the program are shown in light blue, functions are shown in dark blue, and outputs are shown in purple. Detailed information on D-Tracer can be found in Supplementary. Figure created using BioRender.com. B. Structure of d_5_ and d_11_-AA.

To use the program, the user first uploads a peaklist and specifies the types of deuterium labels used in their experiments. The labels can be separated in mass by any number of deuterium atoms. D1 refers to the smaller number of atoms (0 is the smallest possible) and D2 refers to the larger one. The program will then scan through the peaklist and search for all species with a mass difference of the two masses.

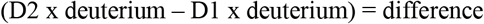

In our case, this value is 6 × 1.0063 or 6.0378 Da. Once a match is identified, the program checks that the retention time is within 0.01 minutes and that the CCS values are within 3% of each other. These tolerance values are hard-coded into the web version but can be manipulated in the source code available on Github. If all three criteria are fulfilled, the feature pair is added to a new dataset. The program will then subtract the mass of the deuterium from the pairs so they can be identified using existing software for non-labeled lipids. To do this, the program subtracts the specified D1 and D2 masses from the smaller and larger mass, respectively, for each pair. The adjusted masses are added as a separate column to the pairs .csv file, which can then be exported for further analysis.

Deuterated lipids were first identified from peak lists containing features that were significantly different between dAA-treated and untreated groups using D-Tracer. The lipids were then validated by manual inspection of the mass spectra. Correctly identified deuterated lipids appear as a doublet or triplet peak on the mass spectra. The doublet peak is formed when d_5_ and d_11_-AA (Figure 2B) are incorporated into the species at a higher concentration than unlabeled AA from the media (Figure 3A). The triplet peak is observed when two AAs are incorporated, giving lipid species with two d_5_-AA, d_5_-AA + d_11_-AA, and two d_11_-AA, separated by 6 Da each (Figure 3A bottom panel). The *m/z* values of the non-labeled version of such lipid species are then calculated manually accordingly.

**Figure 3.**
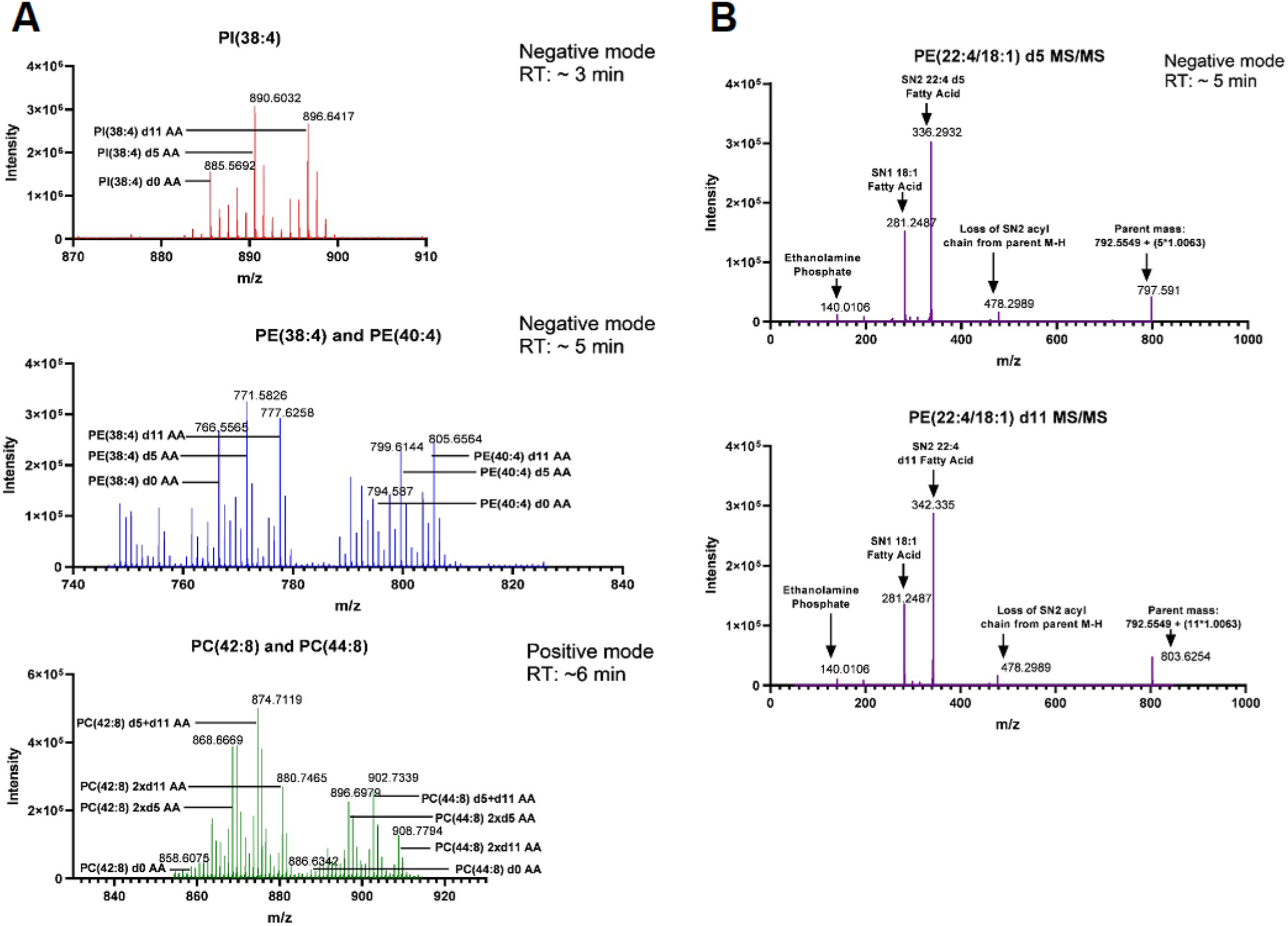
A) Examples of signature doublet and triplet peaks from lipids that incorporate d_5_ and d_11_ AA. Peaks labeled d0-AA represent the lipid incorporating unlabeled AA. PC 42:8 and 44:8 each incorporate two AA, so they form triplets corresponding to 2 x d_5_ AA, 1 d_5_ and 1 d_11_ AA, and 2 x d_11_ AA. B) Targeted MS/MS fragmentation of PE(22:4/18:0) incorporating d_5_ AA and d_11_ AA, respectively. The 22:4 fatty acid is an elongation product of AA 20:4.

Once dAA-containing lipids were extracted from the peaklists with D-Tracer, the resulting mass-adjusted lipid species were processed further using *Lipydomics*, which identifies lipids based on RT, CCS, and *m/z* values.^35^ The identifications were cross-validated using Lipid MAPS MS bulk data search, which searches for *m/z* matches in its lipid database.

### Dual-Isotope Labeling Reveals Differential Incorporation of Exogenous AAs into Various Lipid Classes

Using the workflow described above, we carried out comprehensive dAA-tracing in the cancer cell line, HT-1080, in the presence of vehicle (Control or C), the ferroptosis inducer RSL3 (R), dAA only (D), or co-treatment with dAA + RSL3 (DR). HT-1080 is a well-established cell line that is susceptible to ferroptosis when treated with various ferroptosis inducers, including RSL3.^9^ We first confirmed increased lipid peroxidation using flow cytometry after treatment with C11-BODIPY,^44,45^ which specifically reacts with peroxyl radicals, suggesting the occurrence of ferroptosis when the cells were co-treated with dAA and RSL3 (see Figure S1). After analysis using D-Tracer and *Lipydomics*, we found that dAAs were incorporated into triglycerides (TG), phosphatidylinositol (PI), phosphatidylethanolamine (PE), phosphatidylserine (PS), and phosphatidylcholine (PC) species. The list of dAA-containing lipids is summarized in Tables S3 and S4 for positive and negative modes, respectively. Representative doublet (one AA chain) and triplet (two AA chains) mass spectra are shown in Figure 3A and S6. Untargeted fragmentation of various lipid classes using MS^E^ (Figure S2-S5) allowed the confirmation of the incorporation of d_5_- or d_11_-AA. Interestingly, elongation of dAA to 22:4 followed by incorporation to phospholipids was also observed as confirmed by targeted fragmentation, supporting the utility of dual-isotope tracking of AA metabolism (Figure 3B).

Quantification of the lipid classes was carried out by comparison with an internal standard from each class with a known concentration (Figure 4). The highest level of dAA incorporation was into TG (69% of total lipid quantity), followed by PI (14% of total lipid). Incorporation into PS, PC, and PE lipids comprised 10% or less of the total quantity in the dAA-only treatment group (Figure 4A). In the dAA+RSL3 treatment group, the level of incorporation into TG decreased from 69% to 64%, while incorporation into PI increased from 14% to 20% (Figure 4B). Incorporation into other classes did not change significantly in the co-treatment group from the dAA-only group. The high degree of incorporation into TG species is likely due to the increased formation of lipid droplets in response to high levels of exogenous fatty acids.^46,47^ The alterations in dAA-containing lipid composition in the presence of RSL3 may be a result of ongoing ferroptosis. For example, TGs may be preferentially oxidized in lipid droplets while PIs may not readily undergo lipid peroxidation. However, the elucidation of the detailed roles of these lipids in ferroptosis may involve quantitation of these lipids in specific organelles.

**Figure 4.**
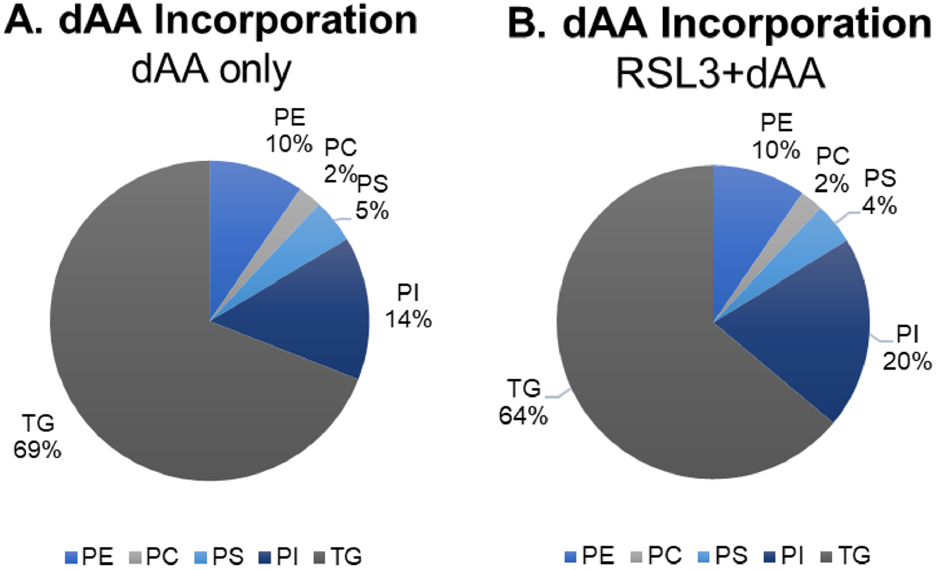
Percent of dAA-incorporating lipids by classes dAA only treatment group (A) and the dAA + RSL3 treatment group (B). The quantity of lipid incorporation was determined by comparison with an internal standard of known concentration. The quantity of each class was divided by the total to determine the percent of each class.

Not all lipid classes incorporated dAA in detectable quantities. Phosphatidic acid (PA) and diacylglycerol (DG) are precursors for PI, PE, PC, and PS phospholipids (Figure 5), but were not found to incorporate dAA. In addition, dAAs were not found in sphingolipids, suggesting sphingolipids do not readily incorporate AAs or promote ferroptosis. PUFA-containing ether lipids have been previously found to promote ferroptosis susceptibility.^48^ Indeed, we found that dAA was incorporated into plasmanyl ether lipids, including PE, PS, and PC ether lipids. The individual species that incorporated dAA and their normalized intensities are shown in a heatmap (Figure 6). As expected, the normalized intensities of all dAA-containing lipids are the highest in dAA-treated groups, D and RD.

**Figure 5.**
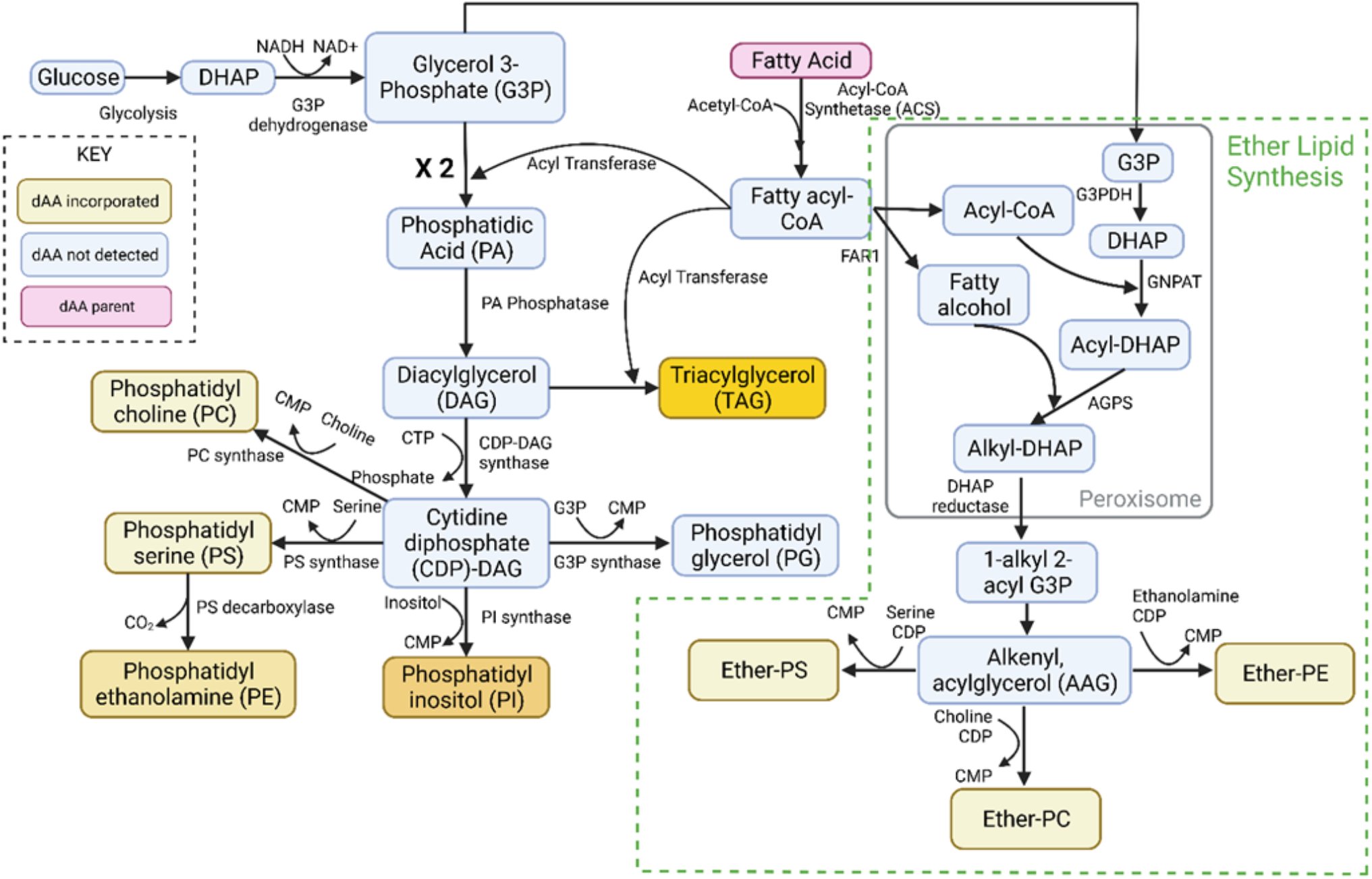
Scheme of the biosynthetic pathway of phospholipids via the cytidine diphosphate diacylglycerol (CDP-DAG) intermediate. Species found to incorporate dAA are shown in yellow. Darker yellow indicates more incorporation, lighter yellow indicates less incorporation as measured by quantitation. Abbreviations: DHAP; dihydroxyacetone phosphate, CMP; cytidine monophosphate, G3PDH; glycerol-3-phosphate dehydrogenase, GNPAT; glyceronephosphate O-acyltransferase, AGPS; alkylglycerone phosphate synthase, FAR1; fatty acyl-CoA reductase. Figure created using BioRender.com

**Figure 6.**
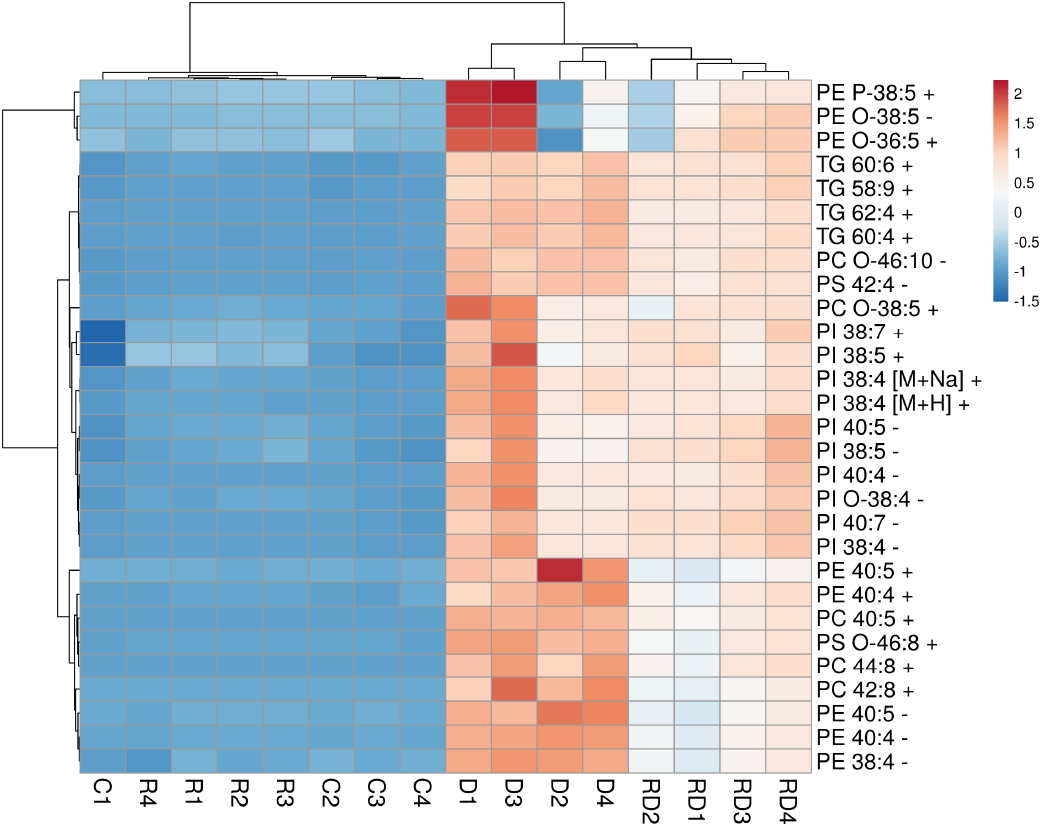
Heatmap of identified dAA-incorporating lipid species and their relative abundances. Rows are centered; unit variance scaling is applied to rows. Rows are clustered using correlation distance and average linkage. Columns are clustered using Euclidean distance and average linkage. Figure created using ClustVis.

In addition, the abundances of lipids in each class, as determined by comparison with the internal standards, were found to be higher in the dAA-only group than in the dAA + RSL3 group although not every class of lipids displayed statistical significance (Figure 7). This could be due to the reduced viability of cells in the co-treatment group, which could lead to overall decreased lipid levels, and/or the oxidation of the parent lipids. Indeed, by manually inspecting the data, we were able to identify an oxidized PE species, PE(38:4) +14Da, which corresponds to +O-2H of PE(38:4) (Figure 8 and S7).

**Figure 7.**
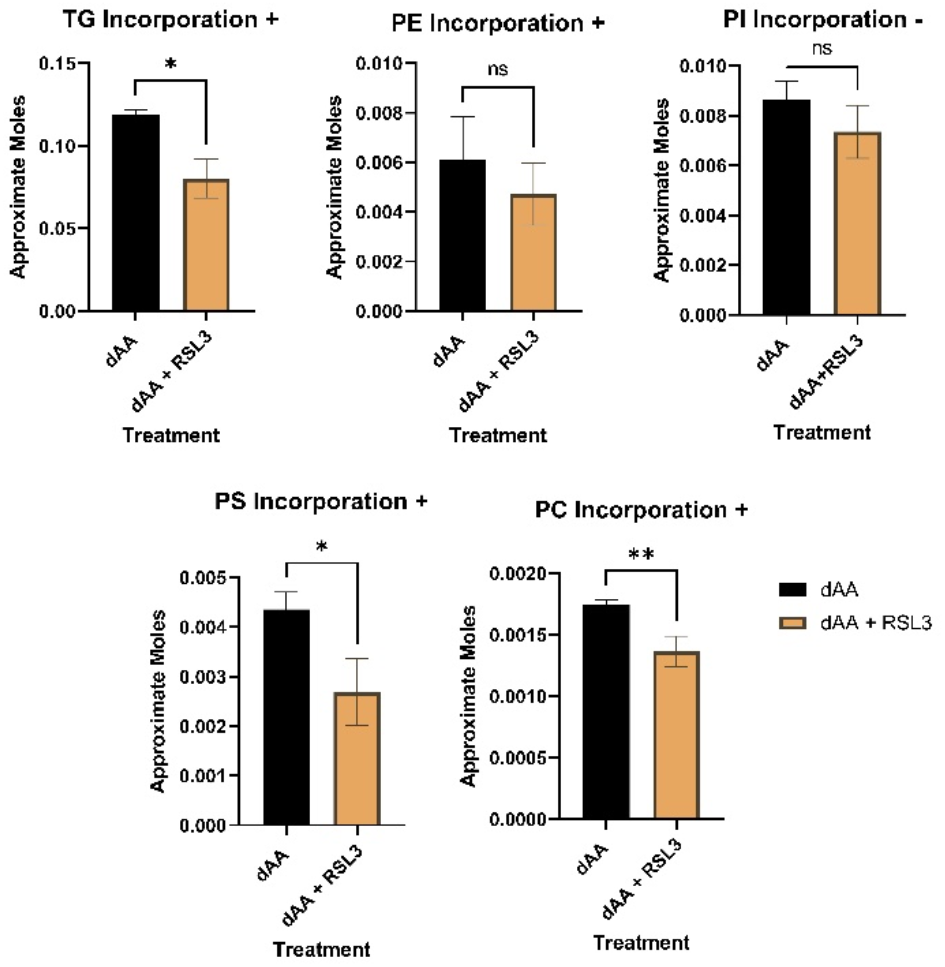
Approximate quantity of dAA incorporating lipids across classes in dAA only and dAA + RSL3 treated cells. Significance was determined by Student’s t-test; *, *P* value < .05. Abbreviations: TG; triglycerides, PE; phosphatidylethanolamine, PI; phosphatidylinositol, PS; phosphatidylserine, PC; phosphatidylcholine. + and – signs in the title indicate that the species was quantified in positive or negative mode, respectively.

**Figure 8.**
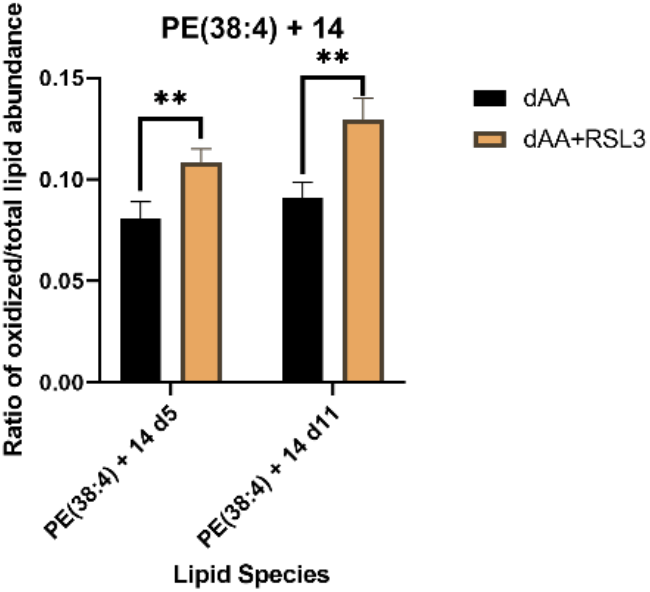
The ratio of the abundance of PE(38:4) + 14 over total abundance of PE(38:4) and PE(38:4) + 14 in dAA only and dAA+RSL3 treatment groups. *P*-values computed using Student’s t-test; **, *P* < 0.002.

Interestingly, the ratio of oxidized lipid : parent lipid was higher in the dAA + RSL3 group than in the dAA-only group, despite the overall lower lipid abundance in the cotreatment group (Figure 8). The identification of the oxidized PE demonstrates the utility of dual-isotope labeling in the confident detection of low-abundance metabolites. The observation of higher level of oxidized PE supports the important roles of PE peroxidation in the execution of ferroptosis, which is consistent with a previous report by Kagan et al.^11^ We note that there are a number of other dAA-containing lipids that were not identified (Tables S3 and S4)through *Lipydomics* and LIPID MAPS. We speculate that these unidentified lipids are also likely oxidized lipids. However, due to the low abundance of these species, targeted fragmentation is not possible to obtain the fatty acid composition. Thus, a limitation of *Lipydomics* is the lack of coverage of the oxidized lipids in its RT and CCS database, and a future direction of this work is to expand the three-dimensional database to include such lipids.

## CONCLUSION

Here we report the development and application of dual-isotope labeling lipidomics to the study of *in vitro* AA metabolism during ferroptosis. We developed a Python software, D-Tracer, to automatically analyze deuterium isotope labeling mass spectra with a user-friendly web interface. Combining this software and *Lipydomics* for lipid identification, we found that deuterated AA is incorporated into several lipid classes in HT-1080 cells, with TG being the predominant one. The abundances of non-oxidized lipids were lower in dAA+RSL3 treated cells relative to dAA treated cells, while the oxidized lipids display an opposite trend, supporting the role of lipid peroxidation in ferroptosis. Thus, the dual isotope labeling combined with three-dimensional HILIC-IM-MS can effectively elucidate lipid metabolic changes associated with biological processes and disease states. The D-Tracer program can also be easily modified to characterize the metabolism of other biological molecules, such as sugar and peptides, potentially expanding its utility.

## Supporting information

Supplemental info-instruction on D-Tracer

## ASSOCIATED CONTENT

### Supporting Information

The Supporting Information is available free of charge on the ACS Publications website.

List of lipid standards, MS parameters, flow cytometry data, untargeted fragmentation mass spectra, examples of mass spectra of dAA-labeled peaks, mass spectra of oxidized lipids, and instructions on using D-Tracer software (separate PDF).

List of dAA-containing lipids in positive and negative modes (Excel files)

## AUTHOR INFORMATION

### Author Contributions

LX conceptualized the study; NR and LX designed the experiments; NR carried out most of the experiments and analyzed the data; NR, RO, AS, and CV developed D-Tracer; AG contributed to cell culture; QD and RZ contributed to lipidomics analysis; NR drafted the manuscript; LX revised the manuscript; all authors have given approval to the submitted version of the manuscript.

## ACKNOWLEDGMENT

We thank the financial support from the National Science Foundation (CHE-1664851) and the National Institutes of Health (R01HD092659).

