## Supplemental tables S1-S2 and figures S1-S7 for "Tracking the Metabolic Fate of Exogenous Arachidonic Acid in Ferroptosis Using Dual-Isotope Labeling Lipidomics"

**Table of Contents:**

1. Table S1: Lipid Internal Standards
2. Table S2: MS parameters used for data acquisition.
3. Figure S1: Lipid Peroxidation by C-11 BODIPY
4. Figure S2-S5: Untargeted Fragmentation Spectra
5. Figure S6: Labeled Peaks Mass Spectra
6. Figure S7: PE(38:4)+14 Mass Spectra
7. Tables S3 and S4: Peaklists and Identifications of dAA containing lipids (Excel files)
8. Supporting Information: D-Tracer (separate .pdf document)

Table S1: Standards included in the lipid internal standard mix.

| Internal Standard | Lipid Class | Vendor | Catalog no. | MW(g/mol) |
|---|---|---|---|---|
| PE (15:0/15:0) | Phosphatidylethanolamine | Avanti | 850704X | 663.906 |
| Cer (d18:1/17:0) | Ceramide | Avanti | 860517P | 551.927 |
| PG (15:0/15:0) | Phosphatidylglycerol | Avanti | 840446X | 716.899 |
| PC (15:0/15:0) | Phosphatidylcholine | Avanti | 850350C | 705.986 |
| PA (12:0/12:0) | Phosphatidic acid | Avanti | 840635X | 558.661 |
| PS (12:0/12:0) | Phosphatidylserine | Avanti | 840038X | 645.738 |
| SM (d18:1/17:0) | Sphingomyelin | Avanti | 860585P | 717.055 |
| LysoPC(15:0/0:0) | Lysophosphatidylcholine | Avanti | 855576C | 481.603 |
| LysoPE (13:0/0:0) | Lysophosphatidylethanolamine | Avanti | 110696 | 411.471 |
| GluCer(d18:1/17:0) | Glucosylceramide | Avanti | 860569 | 714.068 |
| d7-PI(15:0/18:1) | Phosphatidylinositol | Avanti | 791641C | 847.116 |
| DG (13:0/13:0) | Diglyceride | NuChek | D-136 | 484.7519 |
| TG (15:0/15:0) | Triglyceride | NuChek | T-145 | 765.2405 |

Table S2: TriWave parameters used for data acquisition on Waters Synapt XS.

| | |
|---|---|
| IMS Wave Velocity | 500 m/s |
| IMS Wave Height | 40.0 V |
| Trap Wave Velocity | 311 m/z |
| Trap Wave Height | 4.0 V |
| Trap Collision Energy | 4.0 eV |
| Transfer Wave Velocity | 380 m/s |
| Transfer Wave Height | 4.0 V |
| Transfer Collision Energy, Function 1 (MS) | 2.0 eV |
| Transfer Collision Energy, Function 2 ($MS^E$) | Low: 35 eV, High: 45 eV |
| Trap Gas Flow | 2 mL/min |
| Helium Cell Gas Flow | 180 mL/min |
| IMS Gas Flow | 90 mL/min |

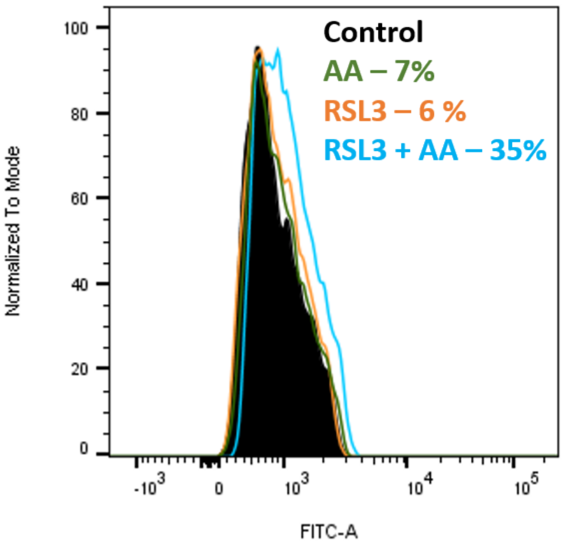


| | Sample Name | Subset Name | Count |
|---|---|---|---|
| | Specimen_001_WT AA.fcs | singlecell | 7920 |
| | Specimen_001_WT RSL3.fcs | singlecell | 8616 |
| | Specimen_001_WT RSL3 + AA.fcs | singlecell | 8357 |
| | Specimen_001_WT Control Stain.fcs | singlecell | 8542 |

Figure S1: Flow cytometry data showing a 35% increase in lipid peroxidation with AA+RSL3 treatment in HT-1080 cells as measured by C11 BODIPY.

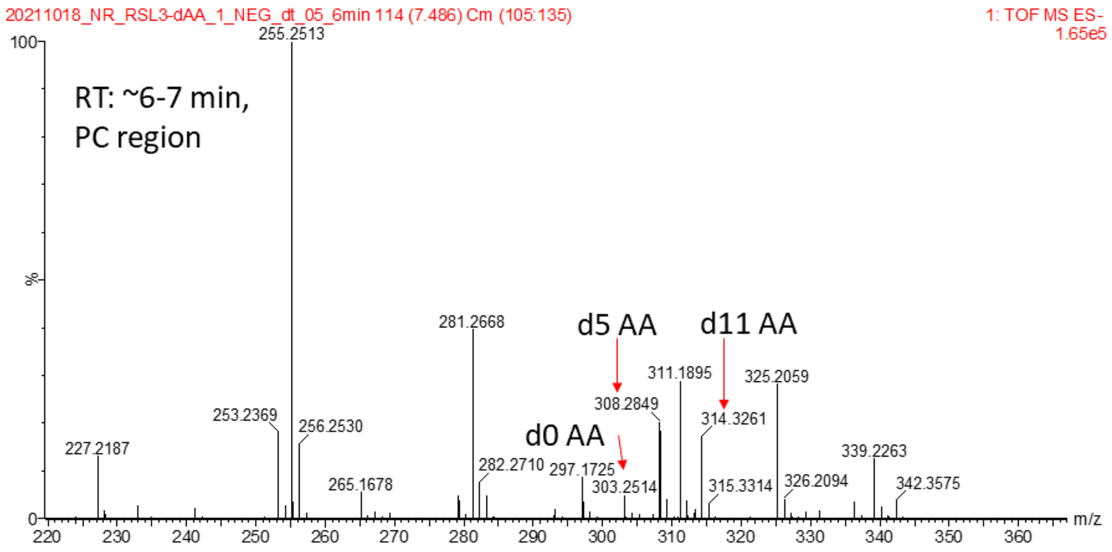


Figure S2: Untargeted fragmentation from the PC, PA, PS region (RT 6-7 min) of the chromatogram. d0 AA, d5 AA, and d11 AA fragments can be seen at m/z 303.2514, 308.2849, and 314.3261, respectively.

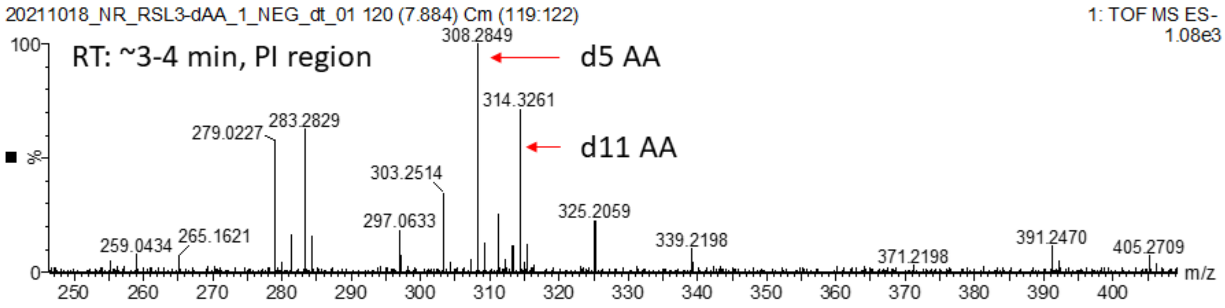


Figure S3: Untargeted fragmentation from the PI region (RT 3-4 min) of the chromatogram. d0 AA, d5 AA, and d11 AA fragments can be seen at m/z 303.2514, 308.2849, and 314.3261, respectively.

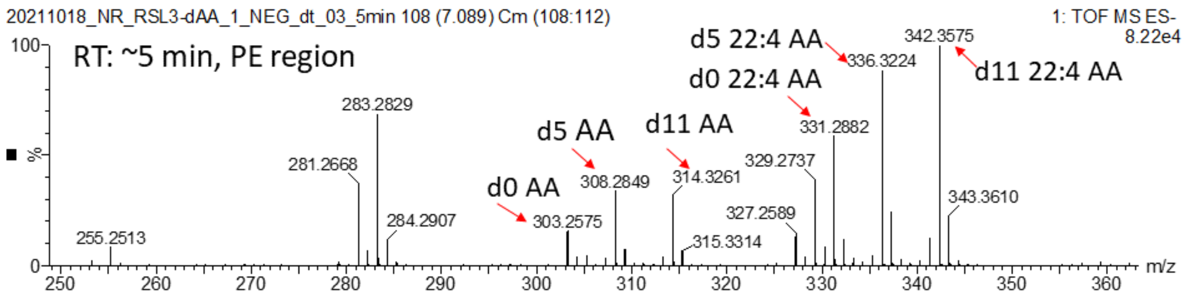


Figure S4: Untargeted fragmentation from the PE region (RT 5 min) of the chromatogram. d0 AA, d5 AA, and d11 AA fragments can be seen at m/z 303.2514, 308.2849, and 314.3261, respectively. d0, d5, and d11 22:4 AA elongation products can also be seen at m/z 331.2882, 336.3224, and 342.3575, respectively.

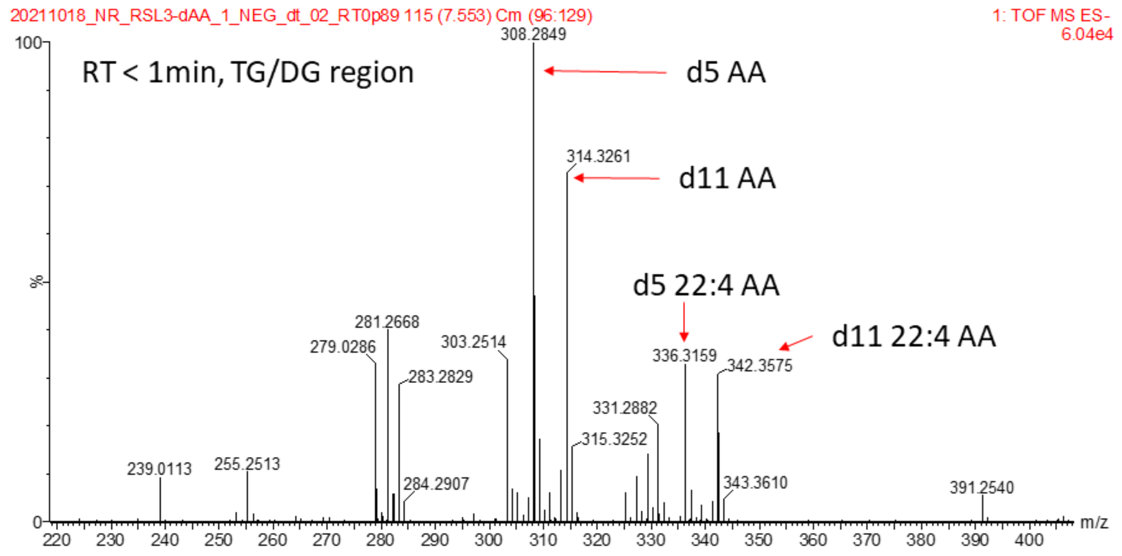


Figure S5: Untargeted fragmentation from the TG/DG region (RT less than 1 min) of the chromatogram. d0 AA, d5 AA, and d11 AA fragments can be seen at m/z 303.2514, 308.2849, and 314.3261, respectively. d0, d5, and d11 22:4 AA elongation products can also be seen at m/z 331.2882, 336.3224, and 342.3575, respectively.

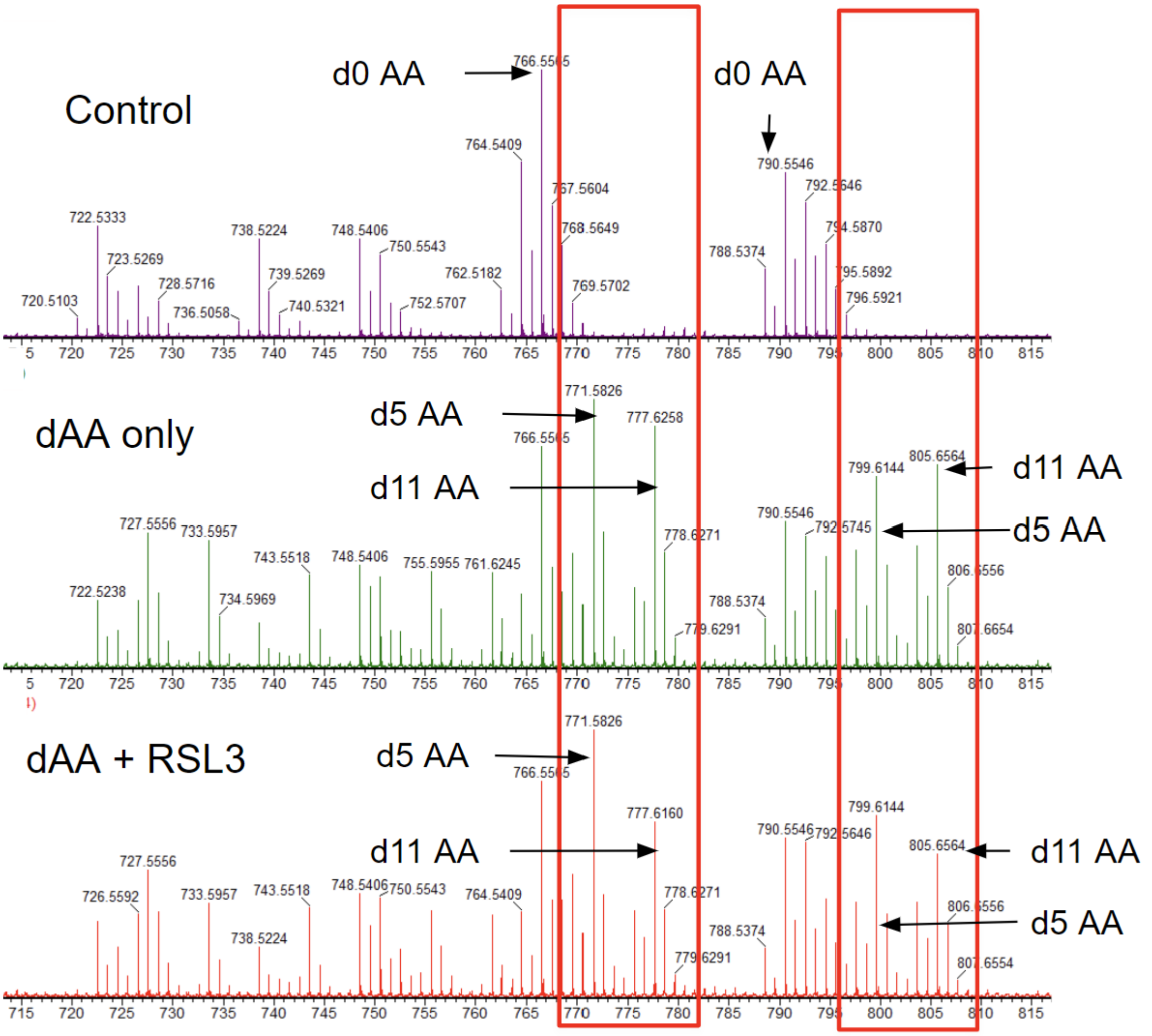


Figure S6: PE region (RT 5 minutes, negative mode) showing two examples of labeled lipids (boxed in red) that are present in dAA treated groups but not in the control group.

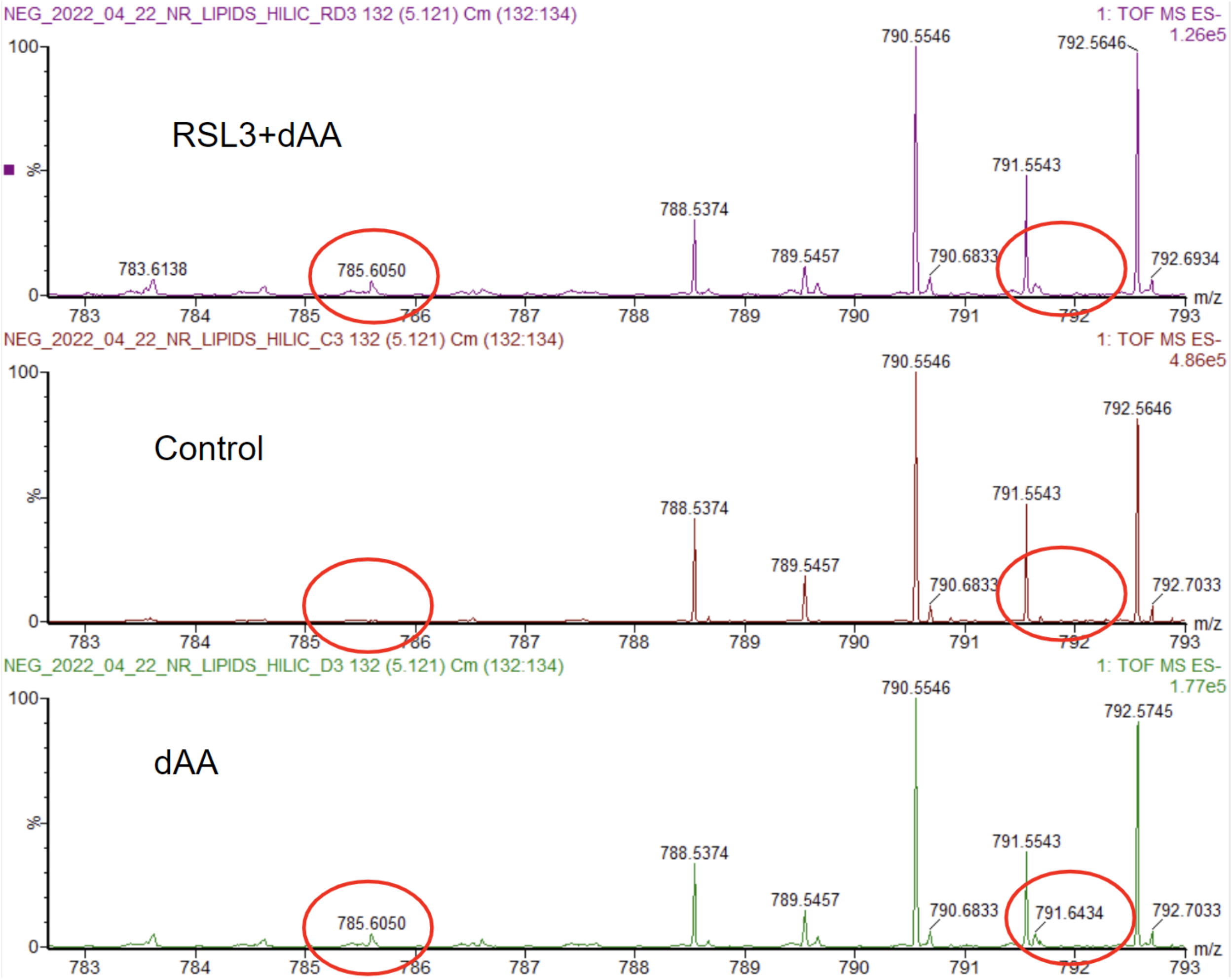


Figure S7: Peaks at 785.6050 and 791.6434 corresponding to PE 38:4 + 14 d5 and d11, respectively in RSL3+dAA, Control, and dAA treated groups. The peaks are only seen in dAA treated groups.
