## Supplemental info-instruction on D-Tracer for "Tracking the Metabolic Fate of Exogenous Arachidonic Acid in Ferroptosis Using Dual-Isotope Labeling Lipidomics"

### **D-Tracer Web Version Instructions**


Noelle Reimers, Ryan Ostrander, Alyson Shoji, Chau Vuong

Contact: | GitHub: nreimers99/d-tracer

University of Washington Department of Medicinal Chemistry


D-Tracer is a program designed for the analysis of dual isotope labeling mass spectrometry data. The user provides a peaklist and D-Tracer will scan through the list to find pairs of compounds with the same retention time and collision cross section values and an m/z difference corresponding to the mass difference of the deuterium labels that were used. It will then adjust the mass of the labeled compounds to the corresponding unlabeled mass so that they can be identified.

User inputs:

- Peaklist.csv file with columns for m/z, RT, CCS, and normalized sample intensities
- Mass of each deuterium label
- Number of samples

Output:

- .csv file containing labeled pairs and isotope-adjusted masses

The web version of D-Tracer can be accessed at this URL:

https://nreimers99-d-tracer-original-d-tracerapplication-xji04u.streamlit.app/

Step 1: Upload dataset. The dataset must be a .csv peaklist containing columns named m/z, Retention time (min), and CCS (angstrom^2). It was originally designed for use with .csv files exported from Waters Progenesis.

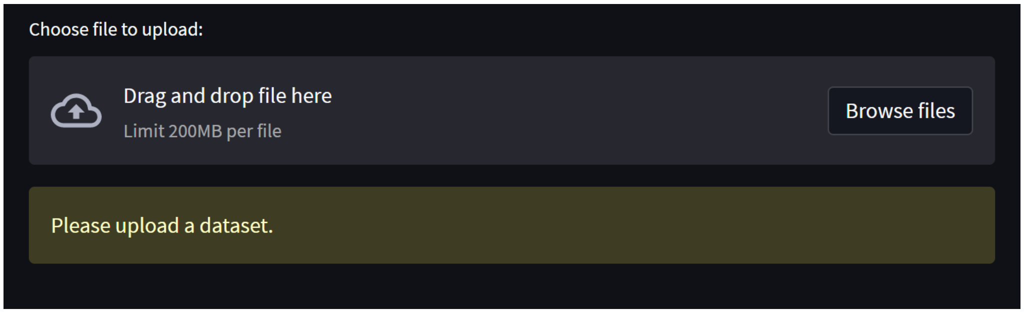


Step 2: The program will accept the file if it has all required columns. The program will raise an error if the file is not adequate for use. Once accepted, the following message will be displayed:

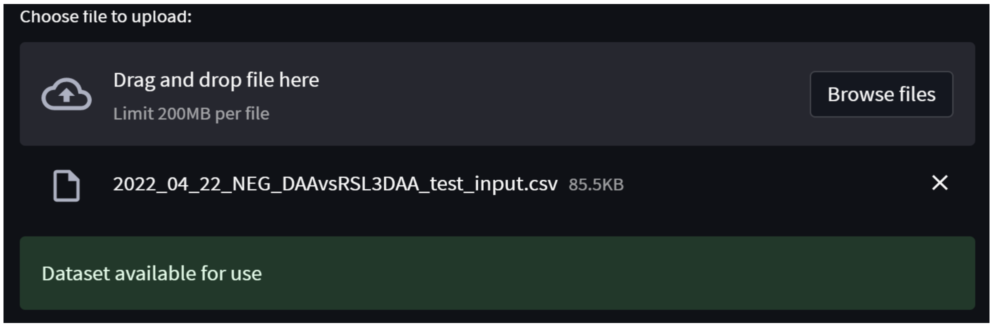


Step 3: Specify parameters. The program needs to know the mass adjustments (ie, the number of deuterium atoms in the two labels) used in the experiment. The first mass adjustment refers to the smaller label used. For example, if the experiment included a label with 5 deuterium atoms and 11 deuterium atoms, the inputs would be 5 for the first mass adjustment and 11 for the second mass adjustment. It also needs to know the number of samples to include intensity data in the output file.

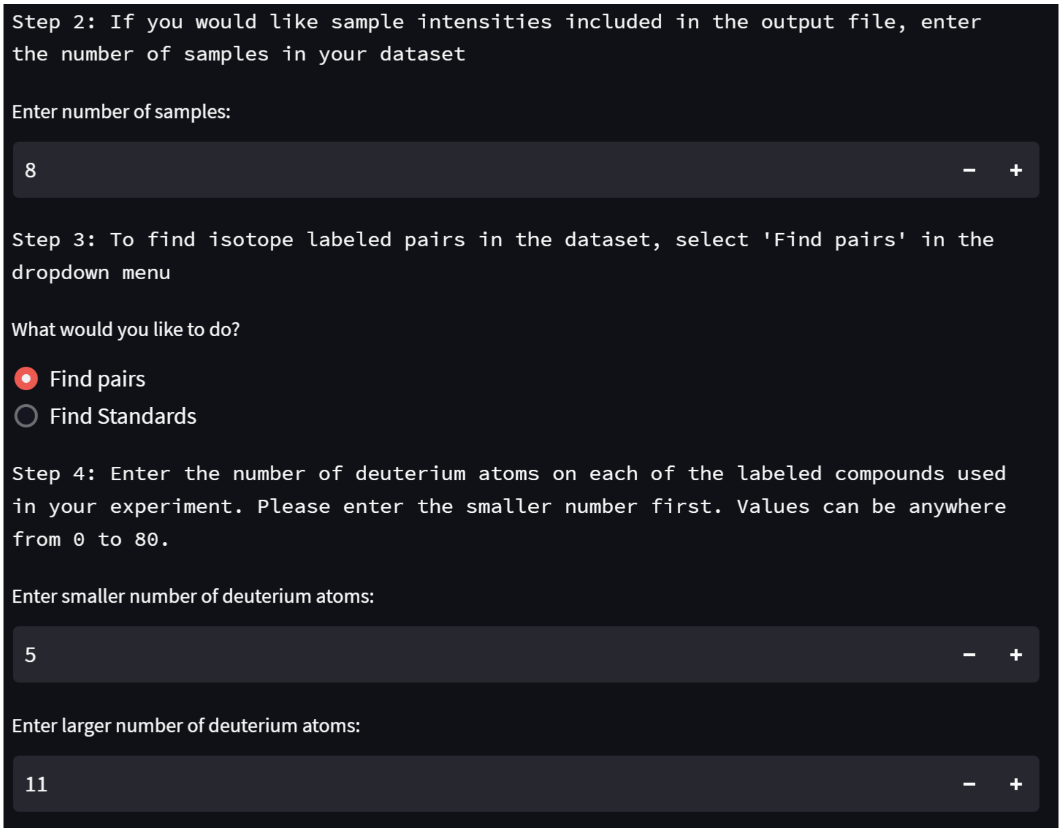


Step 4: Check 'Analyze' to run the pair picking algorithm on the provided dataset. If there are pairs found, the following message will be displayed:

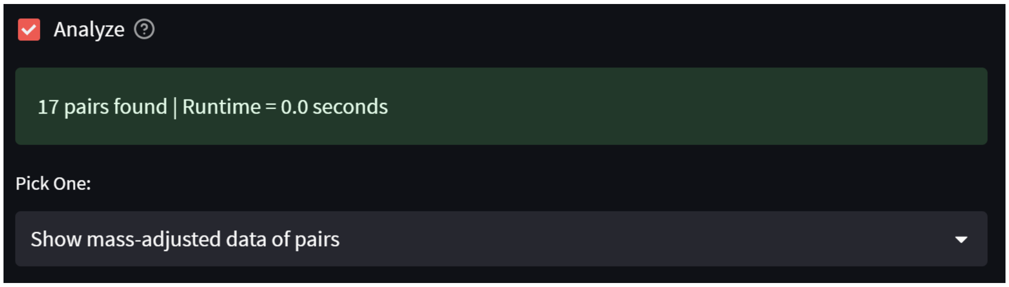


Step 5: Select Show mass-adjusted data of pairs in the dropdown menu to display the pairs found. To download the .csv file to your device, select ‘Export to CSV’ in the bottom left corner.

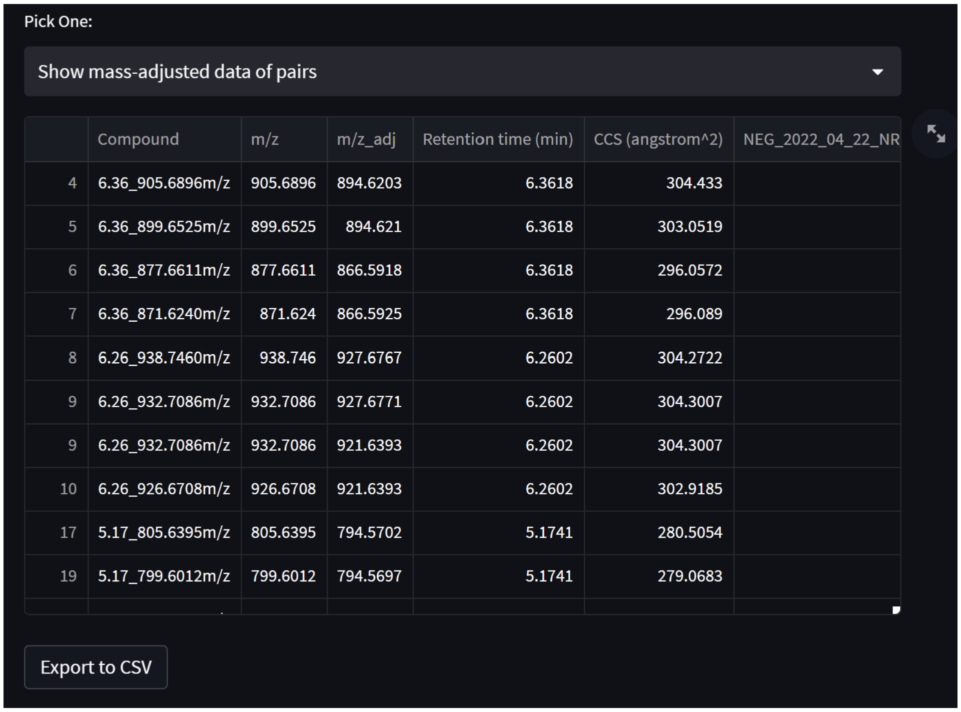


| | Compound | m/z | m/z_adj | Retention time (min) | CCS (angstrom^2) | NEG_2022_04_22_NR |
|---|---|---|---|---|---|---|
| 4 | 6.36_905.6896m/z | 905.6896 | 894.6203 | 6.3618 | 304.433 | |
| 5 | 6.36_899.6525m/z | 899.6525 | 894.621 | 6.3618 | 303.0519 | |
| 6 | 6.36_877.6611m/z | 877.6611 | 866.5918 | 6.3618 | 296.0572 | |
| 7 | 6.36_871.6240m/z | 871.624 | 866.5925 | 6.3618 | 296.089 | |
| 8 | 6.26_938.7460m/z | 938.746 | 927.6767 | 6.2602 | 304.2722 | |
| 9 | 6.26_932.7086m/z | 932.7086 | 927.6771 | 6.2602 | 304.3007 | |
| 9 | 6.26_932.7086m/z | 932.7086 | 921.6393 | 6.2602 | 304.3007 | |
| 10 | 6.26_926.6708m/z | 926.6708 | 921.6393 | 6.2602 | 302.9185 | |
| 17 | 5.17_805.6395m/z | 805.6395 | 794.5702 | 5.1741 | 280.5054 | |
| 19 | 5.17_799.6012m/z | 799.6012 | 794.5697 | 5.1741 | 279.0683 | |

Step 6: If you would like to see a heatmap of intensities for each compound and sample in the dataset, check the ‘Heatmap’ box:

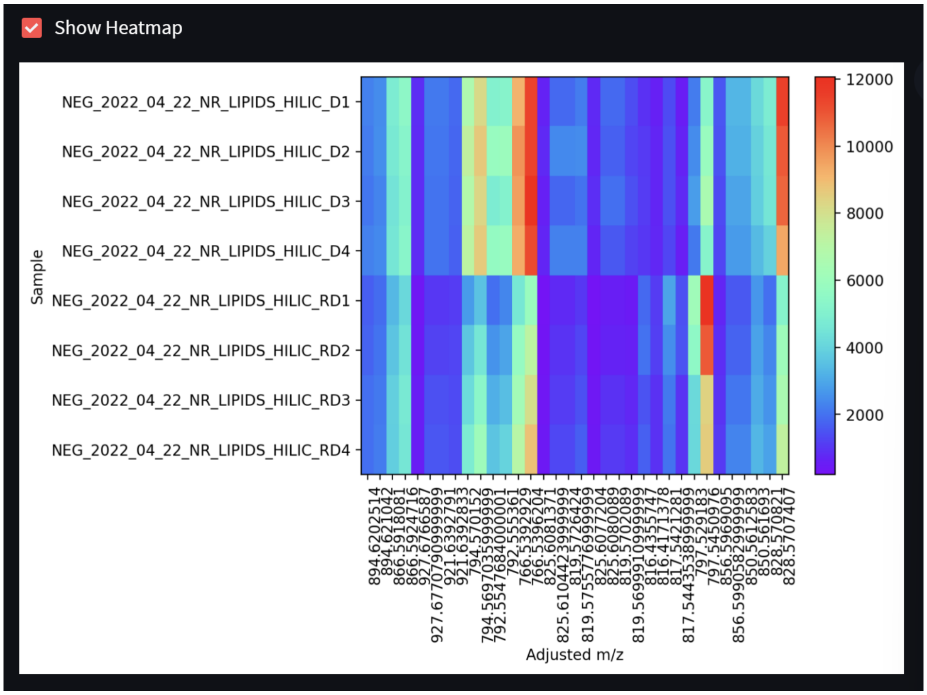
Show Heatmap
Sample
NEG_2022_04_22_NR_LIPIDS_HILIC_D1
NEG_2022_04_22_NR_LIPIDS_HILIC_D2
NEG_2022_04_22_NR_LIPIDS_HILIC_D3
NEG_2022_04_22_NR_LIPIDS_HILIC_D4
NEG_2022_04_22_NR_LIPIDS_HILIC_RD1
NEG_2022_04_22_NR_LIPIDS_HILIC_RD2
NEG_2022_04_22_NR_LIPIDS_HILIC_RD3
NEG_2022_04_22_NR_LIPIDS_HILIC_RD4
12000
10000
8000
6000
4000
2000
Adjusted m/z
